# Identification of the Likely Orthologues of RCD1 within the Plant Family Brassicaceae

**DOI:** 10.1101/2020.04.10.035501

**Authors:** Beena Siddiqua, Syeda Qamarunnisa, Abid Azhar

## Abstract

RCD1 is a signal transduction factor binding protein that gateways a myriad of developmental and stress-related pathways. It was first reported in the wild plant *A. thaliana. Brassica napus* is a cultivated member of the family Brassicaceae, in which the presence of this gene was reported. Using the homology data of these two family-related species, gene for this protein was mined within the genomes of *Brassica carinata, Brassica juncea* and *Brassica oleracea*, using sets of degenerate primers designed on homologous portions of the *A. thaliana* and *B. napus* orthologues. The newly identified sequences were then compared and studied using *in-silico* means and their 3D structures were modelled for having an estimate on their functions. Results demonstrate intergeneric conservation of this protein’s domains on structural and functional levels. The newly found orthologues show potential to be regulated under salinity and oxidative stresses apart from being involved in several developmental stages. These homologues are in-stable *in-vivo* and bear motifs for binding a wide-variety of transcription factors. The structure superimposition studies suggest that these Brassica orthologues bear the WWE domains having transferase activity, the fact that can dramatically increase the survival of these agriculturally important crop plants amid the adverse environmental conditions.

## INTRODUCTION

Genes of importance for the survival of plants are retained within their genomes as a function of selective evolution. Obsolete deletions or complete absence of such important genes make it difficult for plants to thrive (Darwin and Bynum 2009). Therefore, the regions of evolutionary importance are found conserved both at genetic and genomic levels, unless the pressures of artificial selection have acted otherwise (Prakash 2000). These genomic regions encode transcription factors, transcription factor binding elements and comprise constitutive coding regions. Upon perception of a change in external environment; such as an increased heat, cold, salinity, drought or pathogen attack; through the generated ROS (Reactive Oxygen Species) the plants’ internal systems are activated for synergistic or antagonistic reactions (Rejeb et al. 2014). The ultimate outcome of such reactions is gene activation or suppression, leading to the transcriptional reprogramming of cells (Mur et al. 2006). An important player in such pathways is RCD1 (Radical-induced Cell Death 1) that has been implicated in both abiotic as well as biotic stress responses (Brosché et al. 2014). Apart from its role in stress responses, it has also been demonstrated to be important for the proper development of a plant (Jaspers et al. 2009; Teotia and Lamb, 2009). Presently understood function of its protein product is of a transcriptional activator that responds to the elevated levels of certain hormones (Salicylic acid, Methyl Jasmonates, etc) to bring about responses such as shade avoidance and transcriptional reprogramming of cells, while its basal levels are implicated in acting as suppressors of gene expression (Overmyer et el. 2000; Ahlfors et al. 2004; Brosché et al. 2014; Wirthmueller et al. 2017). The RCD1 protein accomplishes these goals by primarily binding to a large array of transcription factors through its N-terminal RST domain (Katiyar-Agarwal et al. 2006; Jaspers et al. 2009; You et al. 2014). And this binding is implicated in regulating the transcriptional control of more than 500 differentially regulated genes (Jaspers et al. 2009).

Since the first report of its presence within *A. thaliana*, the occurrence of this gene has been reported from *Selagenella moelendorfii* to all presently explored land plants’ genomes (Belles-Boix et al. 2000, You et al. 2014). However, its extensive characterization has been carried out in only in the model plant of *A. thaliana*. Where it has been found to be the most active member of SRO family. The members of this family harbour a central PARP domain and an N-terminal RST domain, while only RCD1 and its closest paralogue SRO1 bear an additional domain named WWE (Ahlfors et al. 2004; Jaspers et al. 2010). The SRO members with PARP-RST structures have been reported only in plant family of Brassicaceae (Jaspers et al. 2010) while another category of such orthologues is LROs that harbour WWE and RST domains. The LROs have yet been reported only in case of the plant family Fabaceae (Siddiqua et al. 2016).

Family Brassicaceae houses several important food crops called Caoles; that include *Brassica juncea, B. napus, B. oleraceae* and *B. carinata*. Varieties of these plants are cultivated for obtaining oil and vegetables. In 2010, 76 million tons of vegetables belonging to this genus were produced with net worth of 14.85 billion USD (http://faostat.fao.org/). Existence of RCD1 within members of Brassica genus opens area for manipulation of this gene for the modulation of stress tolerance characteristics of these cultivated plants. From this genus, studies on RCD1 homologue from *B. napus* has been carried out (Anjum et al. 2015). Here we report the existence of members of the WWE bearing SRO family from *B. oleracea, B. carinata* and *B. juncea*; and make a comparative assessment of their putative products using *in-silico* means.

## METHODS

### Obtaining sequences of orthologues

DNA was extracted from *B. carinata* (16195-NARC), *B. juncea* (1664-NARC) and *B. oleracea* (1739-NARC) and *B. napus* (1679-NARC) and *A. thaliana* (Col-0) using standard protocol (Doyle and Doyle, 1987). To these DNA, primers applied were designed using the PRIMER3 software (Untergrasser et al., 2012). The primers were designed for three conserved regions of *A. thaliana* (TAIR Accession: AT1G32230.1) and *B. napus* (NCBI accession: BQ858405) orthologues of *rcd*1, i.e. upstream-WWE (seq1), WWE-PARP (seq2), PARP-RST (seq3), such that these fragments had overlaps at their 3’ junctions. These designed primers (enlisted in Table I) were applied to the extracted DNA. In these reactions, *B. napus* was used as the positive control. PCR was performed in triplicate and repeated at least thrice for the seq1, seq2 and seq3 of each individual species using the PCR recipe: 1X PCR buffer, 1.5Mm MgCl_2_, 0.2Mm dNTPs, 0.5 μM of each primer and 1.5 U of Taq Polymerase. Initial denaturation was achieved at 94°C for 4 mins, 35 cycles of denaturation at 94°C for 50 seconds, annealing at 54°C for 1 minute 30 seconds, extension at 72°C for 1 min while final extension for 10 mins at 72°C. The amplified fragments were purified using kit (MOLEQULE-ON, New Zealand) and amplified using cloning vector pTZ57R/T using InsTAclone PCR Cloning Kit (Thermoscientific, USA) and IPTG/Xgal selection. Successfully amplified fragments were checked by colony PCR and their sequences were determined by sending to a commercial laboratory (MOLEQULE-ON, New Zealand).

**Table 1:**
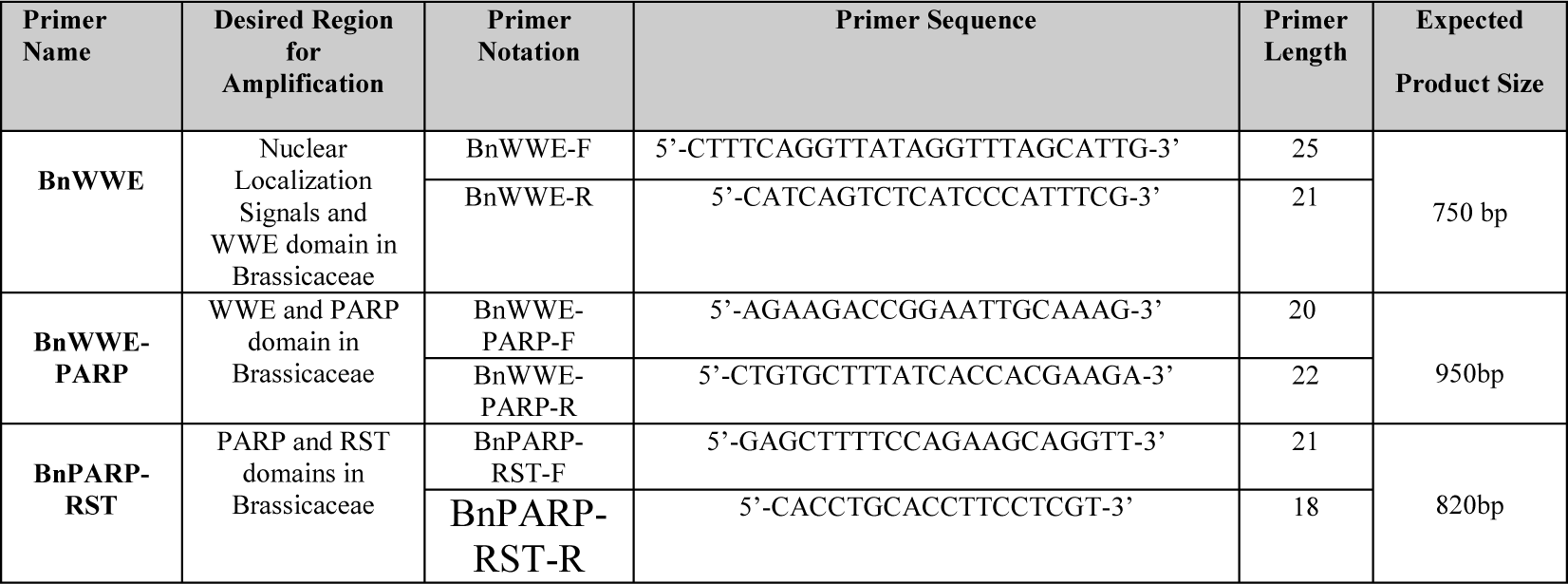
Primers used in Amplification of the *rcd*1 Reads

### *In-silico* analyses of the orthologues

Reads designated as seq1, seq2 and seq3 for each individual species were aligned using the MULTALIN server (Corpet 1988) to obtain full-length orthologue of the *rcd*1 for each individual species. Their reverse sequences were determined using the Reverse Compliment Finder (Stothard 2000), ‘Open Reading Frames’ were determined using the ORF Finder (Rombel et al. 2002), conceptually translated using the ‘Expasy Translate Tool’ (Gasteiger et al. 2003), while their putative domains were identified using ‘InterPro’ (Finn et al. 2016), similarity searches were performed using BLASTp against the refseq_proteins database (Altschul et al. 1997), physical parameters were estimated using “PROTPARAM” (Gasteiger et al., 2003), while its subcellular signals were defined by using WOLFPSORT (Horton et al. 2007). Conceptually translated versions of the homologues were subjected to the determination of their folding capacities using ‘Foldindex’ (Priluskyet al. 2005). From the translated sequences, protein secondary structural elements were determined using CFSSP tool (Chou and Fasman, 1974), while, QUARK and i-TASSER servers (Yang et al. 2015; Xu and Zhang, 2012) were used for the 3D structure predictions. Full chain structures gave a low confidence score (data not shown where TM<0.5), hence individual domains (WWE, PARP and RST) were used for the structure predictions. Tertiary structure models, corresponding to the orthologues, were verified using verify3D (using residues/3D-ID Scores) (Eisenberg et al. 1997). Probable binding sites within the structures were predicted using COACH metaserver to get an idea on putative ligands (Yang et al. 2013) and the function prediction using *in-silico* models was performed using CO-FACTOR (Zhang et al. 2017). For the assessment of transcription factor binding using CDS of homologues, PlantTFDB 4.0 was used (Jin et al. 2017).

## RESULTS

From the extracted DNA, the designed degenerate primers amplified products of nearly 800bp for seq1, 1200bp for seq2 and 1600bp for seq3 (Figure I). The overlapping sequence was trimmed manually from the 5’-ends of the reads. After removing the sequence overlaps, gene sequences obtained were found to have coding sequences (CDS) of 1664bp in case of *B. carinata*, 1954bp in case of *B. juncea* and 2006bp of *B. oleracea*. These sequences were submitted to NCBI, and their issued accessions are MG570396, MG570397 and MG570398, respectively. Their corresponding translated products were 544 amino acids in *B. carinata*, 552aa in *B. juncea* and 562aa in *B. oleracea*. These *in-silico* translated products were named as: BoRCD1, BcRCD1 and BjRCD1, corresponding to the initials of the *Brassica oleracea, Brassica carinata* and *Brassica juncea*. These proteins were found to bear the conserved domains of the RCD1/SRO1 type proteins, which are an N-terminal WWE (InterPro: IPR004170, ProSite: PS50918), central PARP (IPR012317, PFam: PF00644, PS51059) and C-terminal RST (IPR022003, PF12174). The presence of characteristic domains of the RCD1-type proteins in these translated products gave a clue to the presence of the versatile gene named *rcd*1 within these plants of agricultural importance. Physical parameters associated with these proteins suggest them to have an acidic nature, and hence they are proposed to bear a net negative charge at the normal physiological pH, based on the *in-silico* analysis (Table II). The instability indexes suggested them to be highly instable proteins within the plant cell environment (Table II).

**Table II.** Various Physical Parameters of the Brassica RCD1 Proteins

| RCD1 Homologue | Genbank Accession (Coding Sequence/Protein Sequence) | Number of amino acids | Molecular Weight (Daltons) | Isoelectric Point (pI) | Instability Index | Hydropathicity Grand Average |
| --- | --- | --- | --- | --- | --- | --- |
| BcRCD1 | MG570396.1/<br>AYF57917.1 | 544 | 62899.28 | 5.74 | 48.21 | - 0.501 |
| BoRCD1 | MG570398/<br>AYF57919.1 | 562 | 62727.43 | 6.36 | 42.03 | - 0.485 |
| BjRCD1 | MG570397/<br>AYF57918.1 | 552 | 54743.08 | 5.14 | 40.31 | - 0.415 |

**Fig. 1.**
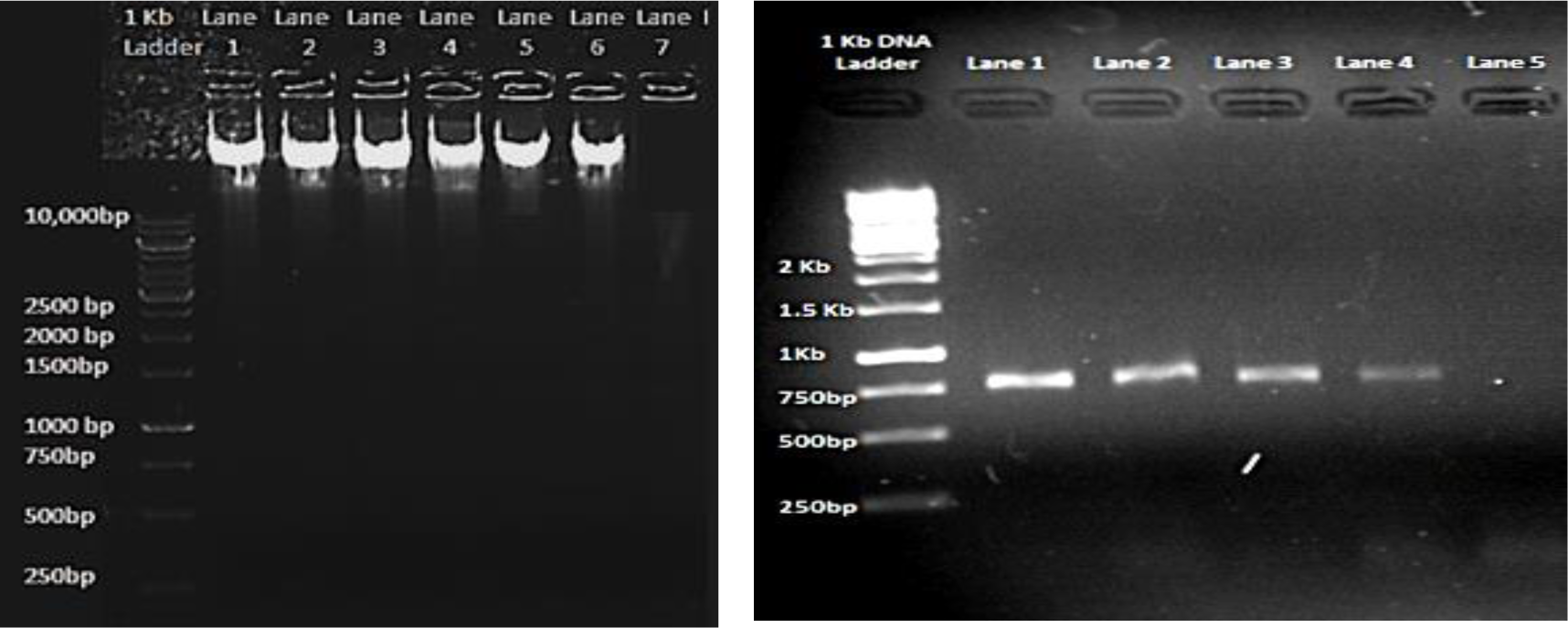
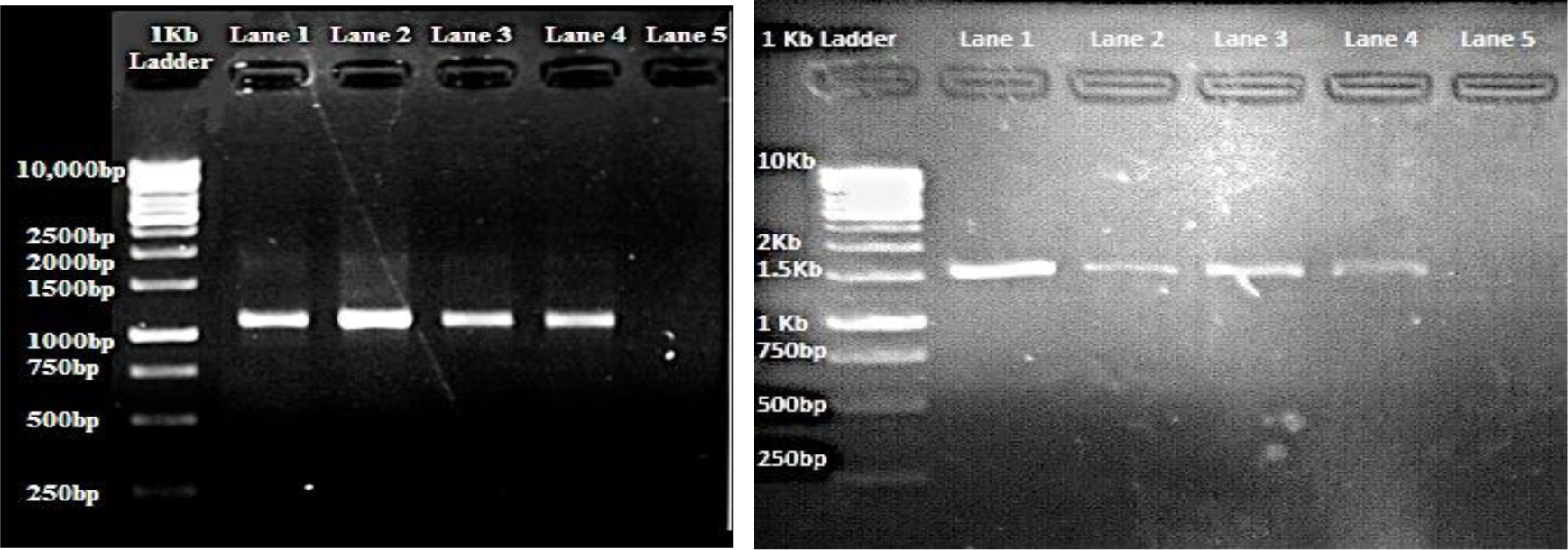
Gel images extracted DNA of *B. carinata* (Lane 1 and 2), *B. juncea* (Lane 3), *B. oleracea* (Lane 4), *B. napus* (Lane 5), *A. thaliana* (Lane 6), Blank (Lane 7). PCR Amplification of the overlapping putative *rcd1* reads with *Brassica oleracea* in lane 1, *Brassica carinata* in lane 2, *Brassica juncea* in lane 3 and *Brassica napus* as positive control in lane 4 of each respective gel. b) Upstream region c) upstream-PARP encoding sequence d) PARP-RST encoding sequence.

Within the subcellular spaces, the BoRCD1, BcRCD1 and BjRCD1 could be localized to several organelles, as indicated by the presence of signals for localization to/across cytoplasm, cytoskeleton, mitochondrion, plasma membrane; in addition to those for nuclear localization (Table III). Out of the three reported NLS (Nuclear Localization Signal) signatures in *A. thaliana* RCD1 (AtRCD1) (by Bellex-bois 2000), KKRKR (NLS1 of AtRCD1) and KKHR (NLS3) were found within the BcRCD1, BoRCD1 and BjRCD1. An important signal of AtRCD1, designated as KRRR (NLS2), was found substituted for KRRKL in the inspected members of the genus Brassica, highlighting an R to K substitution in the signals of these Brassica genus plants.

**Table III.** Prediction of Signals for Subcellular Localization.

| Brassica Homologue | Identifiers For Subcellular Localization |  |  |  |  |
| --- | --- | --- | --- | --- | --- |
|  | Nuclear | Cytoplasmic | Mitochondrial | Plasma Membranic | Cytoskeletal |
| BcRCD1 | At3g09840.1<br>At5g04940.1<br>At5g06460.1, At4g16420.1 | EF2_BETVU<br>At1g77520.1<br>THS1_VITVI<br>At5g37170.1<br>EF2_CHLKE<br>THS2_VITVI | At4g13940.1 | NRL3_ARATH | At2g35630.1 |
| BoRCD1 | At3g09840.1<br>At5g04940.1<br>At5g06460.1 | EF2_BETVU<br>ACOC_ARATH<br>At1g77520.1<br>THS2_VITVI<br>At5g37170.1<br>THS1_VITVI<br>SYK_LYCES | At4g13940.1 | NRL3_ARATH | ACT7_ORYSA<br>At2g35630.1 |
| BjRCD1 | At3g09840.1<br>At5g04940.1 | PPC1_UROPA<br>At1g76990.1<br>PPC2_UROPA | At4g13940.1 | NRL3_ARATH | ACT1_ARATH<br>ACT2_DAUCA<br>ACT13_SOLTU<br>ACT1_SORBI<br>ACT1_ORYSA |

Foldindex analysis revealed that these orthologues have 2 highly folded areas within the PARP region (residues 250 to 350) and a net total of 248-262 disordered residues (Supplementary data Fig I). Presence of disordered region supports the participation of these proteins in regulatory processes that affect a plant’s survival. Prediction of transcription factor binding sites using p-value threshold set at p-value ≤ 1e-5, resulted in prediction of 48 binding sites of 34 transcription factors from BoRCD1. Similar searches identified 28 sites from 26 transcription factors from BcRCD1 and 38 binding sites from 34 TFs from BjRCD1. Binding to an array of transcription factors is an attribute of a regulatory/hub proteins that take part in regulatory processes of a plant (Vandereyken et al. 2018). The homologues bear motifs for interaction with transcription factors belonging to several families (TFs summarised in table IV), importantly with those belonging to AP2/ERF, as 25 such binding motifs have been identified for BoRCD1, 14 BcRCD1 and 12 for BjRCD1.

**Table IV.** Prediction of Transcription Factor Binding of Brassica Homologues

| **Brassica species** | **Transcription factors** |
| --- | --- |
| *B. carinata* | ARF, E2F/DP, ERF, GATA,LBD, MYB, NAC,SBP, Trihelix |
| *B. oleracea* | ARF,<br>Bzip,E2F/DP,ERF,GATA,LBD,MIKC_MADS,MYB,NAC,SBP,Trihelix,<br>WKRY |
| *B. juncea* | ARF,ARR-B, B3,BBR-BPC,C2H2,E2F/DP,ERF,GATA,<br>LBD,MIKC_MADS,MYB,NAC, Trihelix,SBP,WRKY |

The 3D structure remodelling using *in-silico* environment was performed for the whole proteins and this gave low scores owing to the presence of several stretches of disordered residues amid domain forming stretches of amino acids (data not shown). This has also been indicated by the Foldindex analysis (Supplementary data Fig I). To get a better resolution, domain forming stretches of amino acids were subjected to *in-silico* structure determination using QUARK and i-TASSER servers. The structures of RST were determined with TM (Template Modelling) scores in the range of 0.7-0.85 (Table V), and revealed the sole presence of alpha-helices being separated by turns and coils leading to an overall structure shown in Fig III. Similarly, the WWE forms an overall structure comprising a single large alpha-helix being separated by two b-sheets through some turns. This domain also shares structural homology to the pyridoxal kinases that have transferase activity *in-vivo*.

**Table V.** Quality Assessment Results of the *in-silico* Models

| Brassica species | Template Modelling (Tm) Score of Models<br>(Range: 0-1) |  |  | Verify3D<br>(Percent Residues with 3D/ID Scores> 0.2) |  |  |
| --- | --- | --- | --- | --- | --- | --- |
|  | WWE | PARP | RST | WWE | PARP | RST |
| *B. carinata* | 0.6148<br>$\pm 0.0764$ | 0.85±0.08 | 0.7967<br>$\pm 0.0764$ | 98.36% | 86.52% | 93.85% |
| *B. oleracea* | 0.5947<br>$\pm 0.0764$ | 0.69±0.12 | 0.8074<br>$\pm 0.0764$ | 97.84% | 93.61% | 90.23% |
| *B. juncea* | 0.6148<br>$\pm 0.0764$ | 0.67±0.12 | 0.8512<br>$\pm 0.0764$ | 98.36% | 96.40% | 90.77% |

**Fig III.**
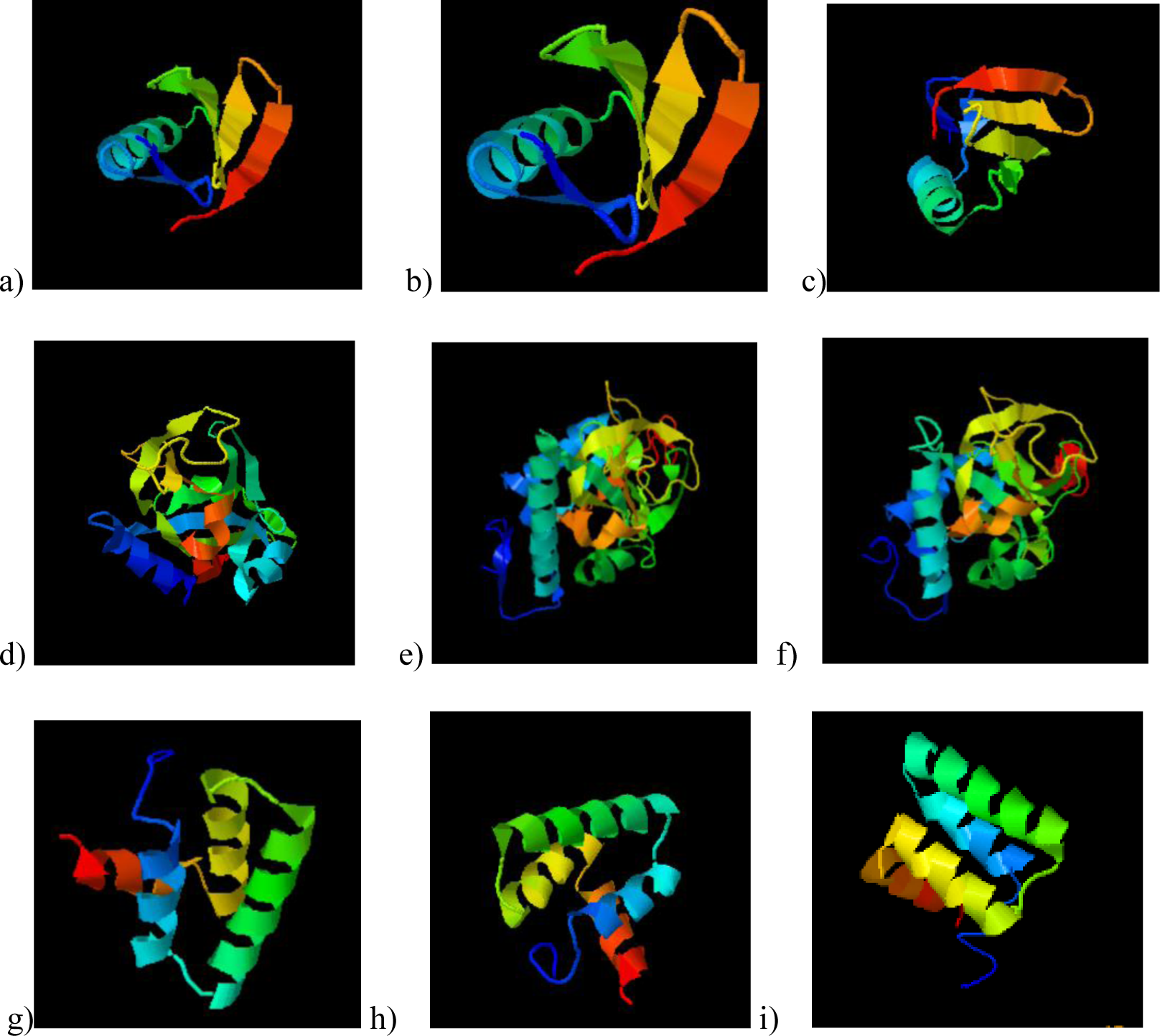
Structures of Brassica homologues’ domains: a) BC WWE b) BJ WWE c) BO WWE d) BC PARP e) BJ PARP f) BO PARP *g)* BC RST h) BJ RST i) BO RST

The structure-based molecular functional prediction for each of the Brassica PARP model was Gene Ontology (GO): 0003950 that outlines structures with an active PARP. The GO scores obtained (out of 1) were 0.90, 0.91 and 0.95 for BjRCD1, BoRCD1 and BcRCD1, respectively. Further studies were made using COFACTOR server, where PARP activity was predicted with a 69% for BcRCD1, 61% for BoRCD1 and 65% for BjRCD1 PARP. In further studies, we found these homologues to have good affinities for binding inhibitors like RGK (2-(4-Aminophenyl)-3,4-Dihydroquinazolin-4-One) and GAB (3-Aminobenzoic Acid) (Fig IV). The Confidence Score (CS) for BcRCD1 PARP with RGK was 0.77, BjRCD1 PARP with GAB was 0.65 and BoRCD1 PARP with GAB was 0.66. These are the inhibitors o PARP activity. The key residues comprising this binding site are predicted to be LP-HLT-FS-Y-N. The PARP structures of newly found orthologues seem to be closely related to the AtRCD1 PARP, as determined using superimposition procedures (Figure IV).

**Fig IV.**
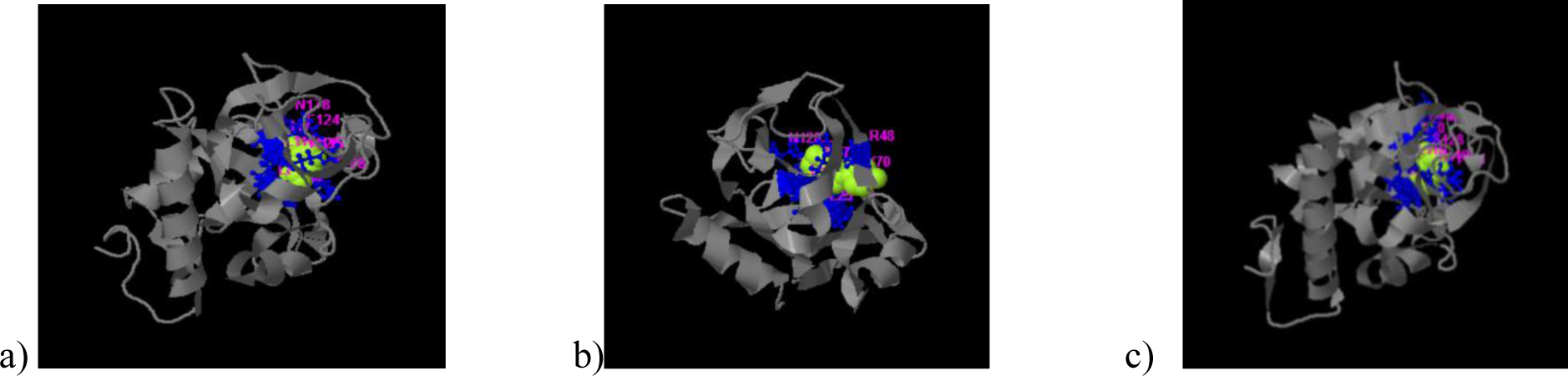
Ligand Binding Prediction of Brassica PARPs a) BC PARP with RGK b) BJ PARP with GAB c) BO PARP with GAB

TM-Align has identified the crystal structure of the RCD1 of *A. thaliana* PARP (with PDB id: 5ngoA) to be closely related to that of *B. carinata* PARP, having an RMSD (Root Mean Square Deviation) of 0.82 over a coverage of 0.986. The TM scores obtained were 0.958/1 for BcRCD1, 0.794/1 for BoRCD1 and 0.779/1 for BjRCD1. The superimposition has been shown in Fig V, indicating these proteins to be closely related in structure and function to the AtRCD1.

**Fig V.**
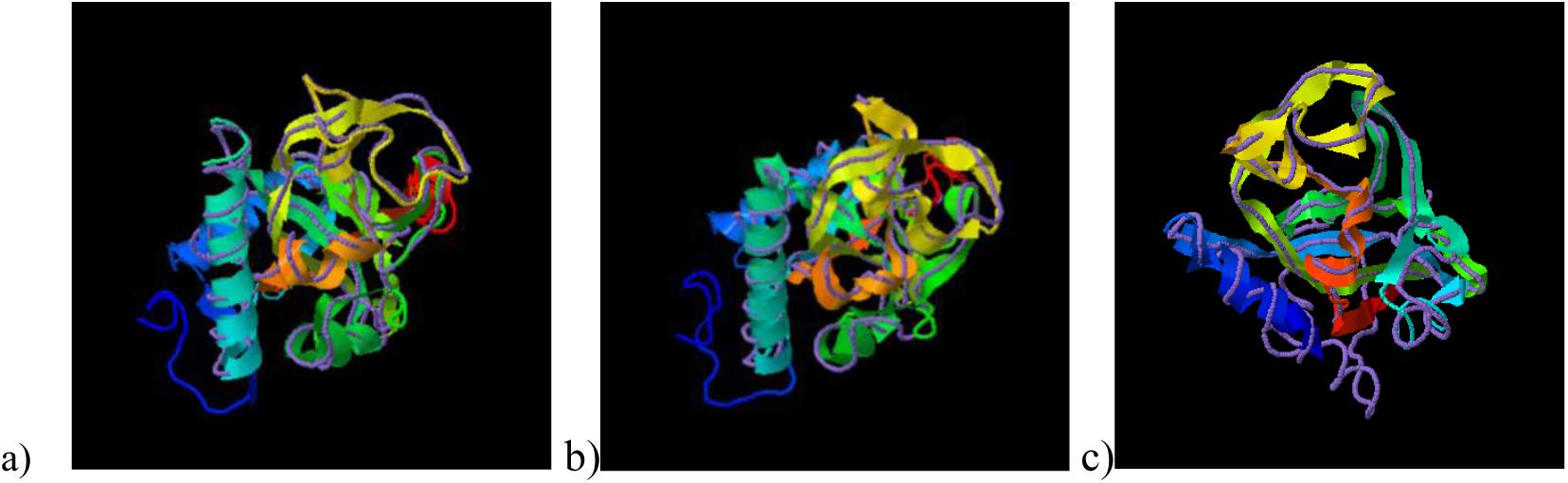
Superimposition of the PARP models of newly found Brassica homologues of RCD1 with the crystal structure of *A. thaliana* PARP (5ngoA) using TM-Align, where ribbon indicates Brassica PARP and discreet line indicates *A. thaliana* PARP. a) *B. carinata* b) *B. juncea* c) *B. oleracea*.

Some greater contribution towards the functions exhibited by these homologues can come from the PARP domain if it’s catalytically active, as indicated in the consensus prediction of the function for PARP (Fig VI); indicating these proteins to be involved in a myriad of developmental and stress-related pathways. Some pathways predicted with a high confidence (>0.9) are response to stress (especially salt stress). Hence, the findings suggest the found genes to encode proteins homologous to the RCD1 of AtRCD1 and highlight a prospect of being a potential candidate for the development of resistant cultivars using breeding and/or biotechnological approaches.

**Fig VI.**
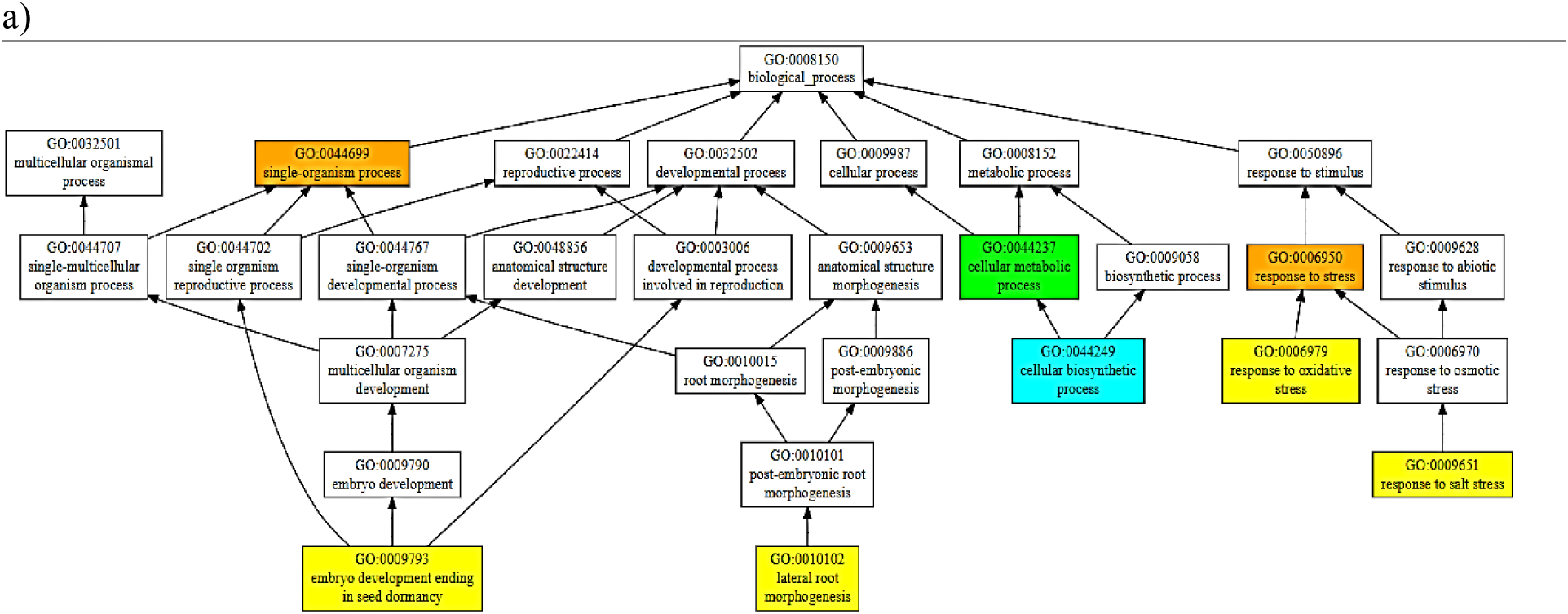
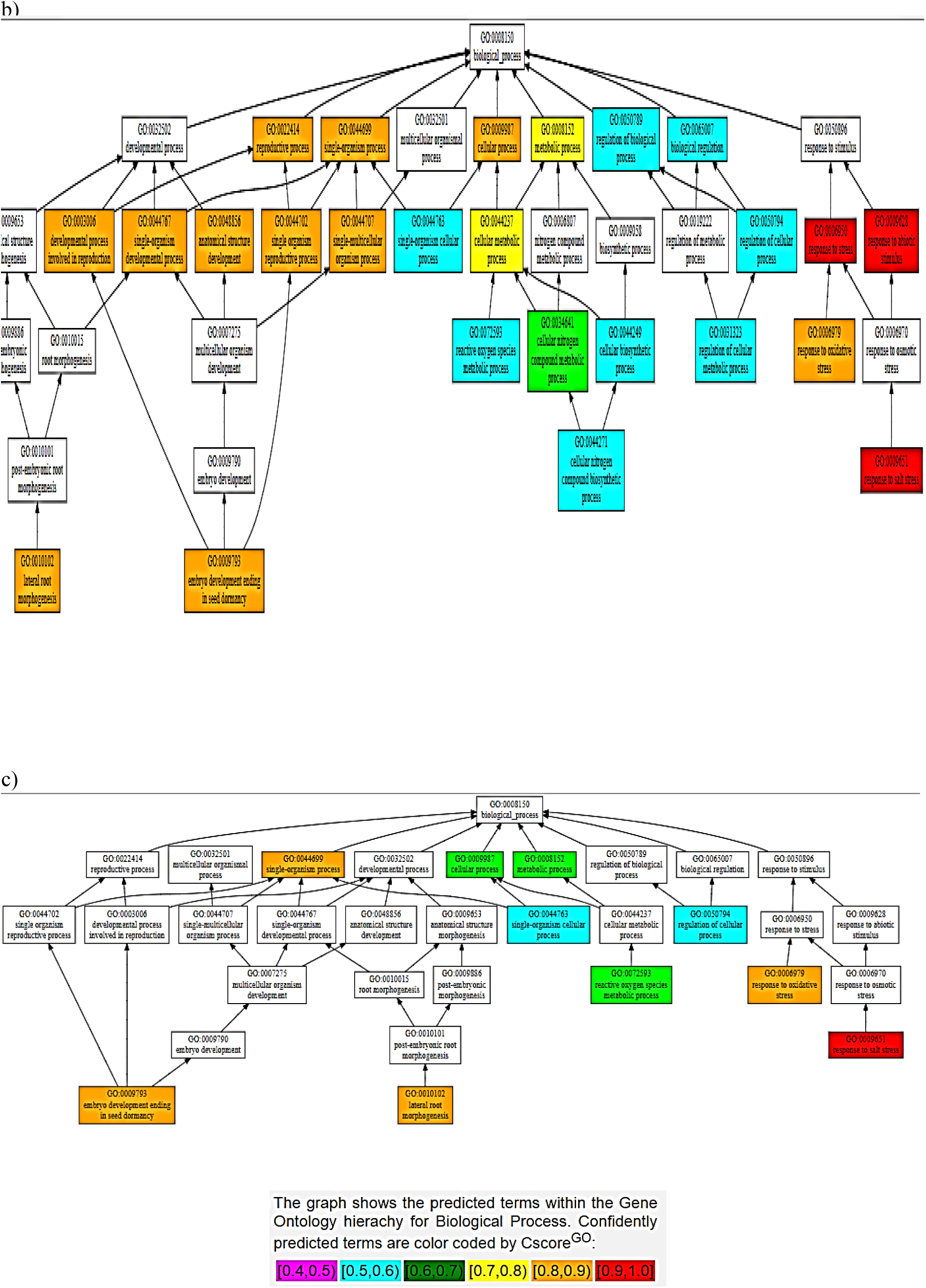
Consensus Prediction of Biological Function of the domains of BcRCD1, BoRCD1 and BjRCD1 a) WWE b) PARP c) RST

## DISCUSSION

Bringing genes that regulate multiple stresses on forefront and studying their properties could play some role in the biotechnological approaches that aim at making crops resilient to the environmental upsets (Meyer and Purugannan, 2013). As part of our current finding, the versatile RCD1 of *Arabidopsis thaliana* has its orthologous relatives within three cultivated varieties of the Brassica genus (*B. carinata, B. juncea, B. oleracea*). The amplification of the orthologues from the family-related species; i.e. *Brassica carinata, B. juncea* and *B. oleracea*, using primers designed on conserved genomic regions from *Brassica napus* and *Arabidopsis thaliana* orthologues of the *rcd*1; demonstrate that these orthologues share some degree of sequence conservation across these closely related plants. This feature has also been demonstrated in a previous research in which the members of various plant families have been found forming phylogenetic clades being supported through high bootstrap values (Siddiqua et al. 2016).

The analysis of the putative protein products encoded by these genes, show that they bear complete WWE domain alongside PARP and RST domains. The WWE domain characteristically exists within the active members RCD1 and SRO1 of *A. thaliana*, both of which demonstrate their functions through genetic redundancies (Jaspers et al. 2009, Teotia and Lamb, 2009). Aside the WWE domain, at N-terminal end of the protein, three signals for nuclear localization, KKRKR (NLS1), KRRR (NLS2), and KKHR (NLS3), have been reported in the AtRCD1 (Belles-Boix et al. 2000). Two of these signals; NLS1 and NLS2, are reported to be essential for the nuclear localization of AtRCD1 under unstressed situations. While upon exposure to stress, AtRCD1 has been shown to be found within cytoplasm as well as in nucleus (Katiyar-Agarwal et al. 2006). These proteins also have instability indices of 40 or above, and such protein are instable in-vivo (Guruprasad et al. 1990). The signals obtained within the conceptually translated Brassica homologues show their import-export into several plant cellular compartments. The absense of the NLS2, which has been shown to be of importance for the nuclear localization of the AtRCD1 by Katiyar-Agarwal and coworkers, indicates some reduced localization of this protein in nucleus in comparison to the AtRCD1. Moreover, the presence of signatures for mitochondrial and chloroplastic import indicate chances of retrograde signalling involving RCD1. Signals for transport to plasma membrane have also been identified within these orthologues of AtRCD1, where SOS1 resides that has previously been indicated to interact with AtRCD1 (Katiyar-Agarwal et al. 2006). The Arabidopsis RCD1 has been shown to regulate signaling from chloroplasts and mitochondria to establish transcriptional control over the regulatory processes in these organelles (Shapiguzov et al., 2019).

Some idea on the functional characteristics of these orthologues was drawn using the *in-silico* structural data. The structure-based function prediction brings important highlights on the *in-vivo* characteristics of a protein (Wilkins et al. 2012). Such approach has revealed these newly found RCD1 homologues to be involved in oxidative and salinity responses among stress-responsive pipeline. This finding corroborates with the previous findings in AtRCD1 that the protein interacts with SOS1 tails and controls set of genes that respond to salt and ROS generated stress signals (Belles-boix et al. 2000; Katiyar-Agarwal et al. 2006). While, among the developmental pathways, the consensus prediction has been of lateral root morphogenesis and embryonic development ending in seed-dormancy. The expression of AtRCD1 in root-tip has been observed using GUS-tagged promotors’ expression (Katiyar-Agarwal et al. 2006). Some similar results were obtained in case of RCD1/SRO1 double mutants in *A. thaliana* when lesser number of seeds sprouted and developed into plants (Jaspers et al. 2009) that gives some clues that the Brassica homologues share functional similarity to AtRCD1. Importantly, this structure-based function prediction has indicated PARP activity, marginally, for these homologues that share some degree of structural analogy to the *A. thaliana* PARP. On the other hand, this *in-silico* finding has also identified PARP domains of BcRCD1, BjRCD1 and BoRCD1 in binding the inhibitors GAB and RGK. This binding to inhibitors could also lead to an inactive PARP. Such suggestion about has also been previously made about AtRCD1 (Jaspers et al. 2010). PARPs regulate highly important cell functions that include gene expression regulation, programmed cell death and DNA damage response, to name a few (Vainonen et al. 2016). Hence, the activity displayed by PARP could modulate the overall function displayed by the RCD1 homologues and could be the potential target of crop-improvement strategies. This is of importance as currently it has been suggested that versatile functions exhibited by AtRCD1 has no involvement of PARP activity (Wirthmueller et al. 2017). On the other hand, it has been demonstrated that the presence of an active PARP within the RCD1 could drastically improve the stress tolerance potential of the reservoir plant (Liu et al. 2014). Hence, owing to the absence of catalytic binding triad of H-Y-E within the binding site and close similarity to the PARP of AtRCD1, it is being suggested that the PARP might not be catalytically active within these orthologues. Based on structure-based function prediction of all domains (Fig 4), we can conclude that these homologues have high chances of being involved in responses to salinity and oxidative stresses among stress-responsive pathways and in embryonic seed development that ends in dormancy and lateral root morphogenesis among the developmental pathways.

The RST domains have important role in binding transcription factors (Katiyar-Agarwal et al. 2006; Jaspers et al. 2009; You et al. 2014). In *A. thaliana*, at least 21 transcription factors have been shown to interact through the RST domains to TFs like DREB2A (of AP2/ERF class) that regulate multiple stresses like drought and heat (Belles-Boix et al. 2000; Jaspers et al. 2009). The TFs predicted to have binding affinities for Brassica homologues fall under diverse classes like MYB, GATA, NAC, that play diverse roles of stress regulation (Reviewed in Lindemoss et al. 2013). Of particular importance are the enrichment of motifs within newly found orthologues for the AP2/ERF class of TFs, as these TFs participate in wide variety of stress and developmental responses and are current targets for crop improvement perspectives (Phukan et al. 2017). In the Brassica orthologues of AtRCD1, the RST domain was found comprising a structure that was solely represented by alpha helices, and this has also been shown to be the case with *A. thaliana* RST whose structural constrains have been determined using NMR (Nucleic Magnetic Resonance) (Tossavainen et al. 2017). Hence, the overall analysis suggests that there exists some degree of conservation with respect to sequence, structure and function across RCD1 orthologues within the family of Brassicaceae.

## Supporting information

Figure 1: Folded and Unfolded areas within the Brassica Homologues of RCD1 a) B. carinata, b) B. juncea c) B. oleracea

---

**Fig 1.**
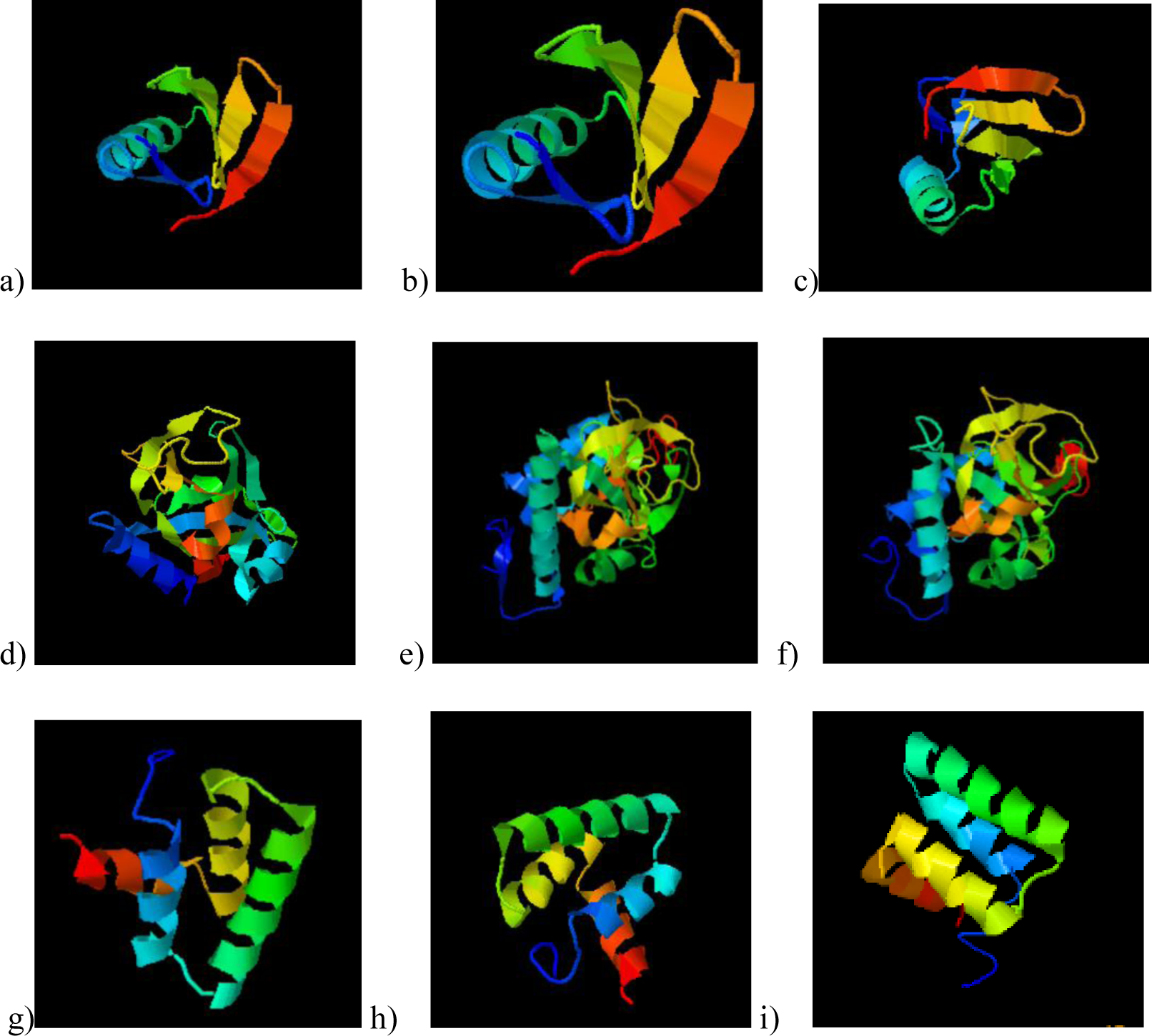
Structures of Brassica homologues’ domains: a) BC WWE b) BJ WWE c) BO WWE d) BC PARP e) BJ PARP f) BO PARP *g)* BC RST h) BJ RST i) BO RST

**Fig 2.**
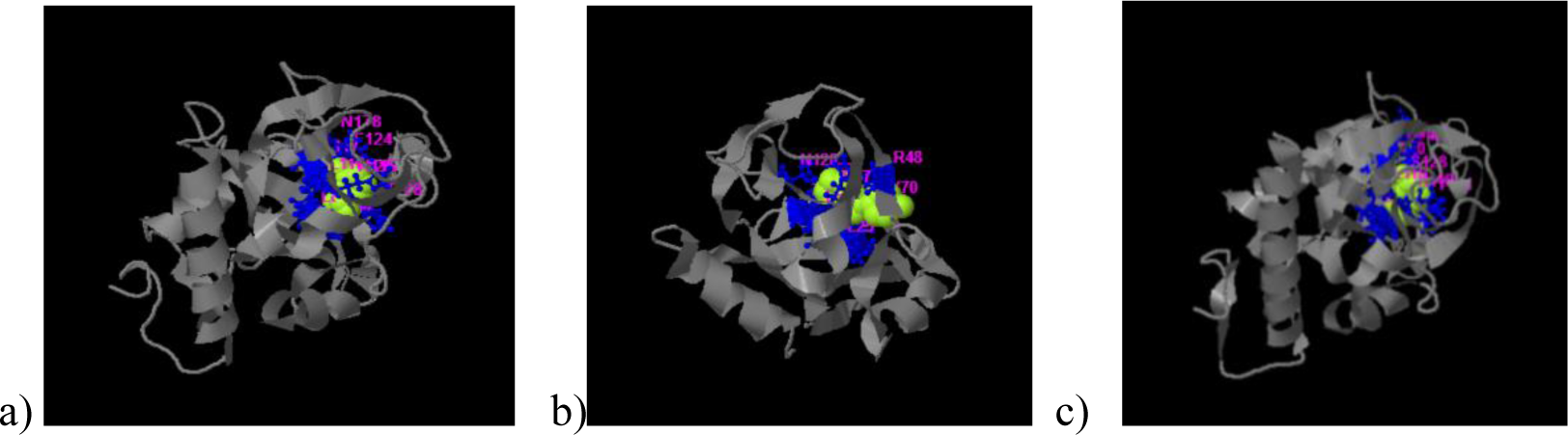
Ligand Binding Prediction of Brassica PARPs a) BC PARP with RGK b) BJ PARP with GAB c) BO PARP with GAB

**Fig 3.**
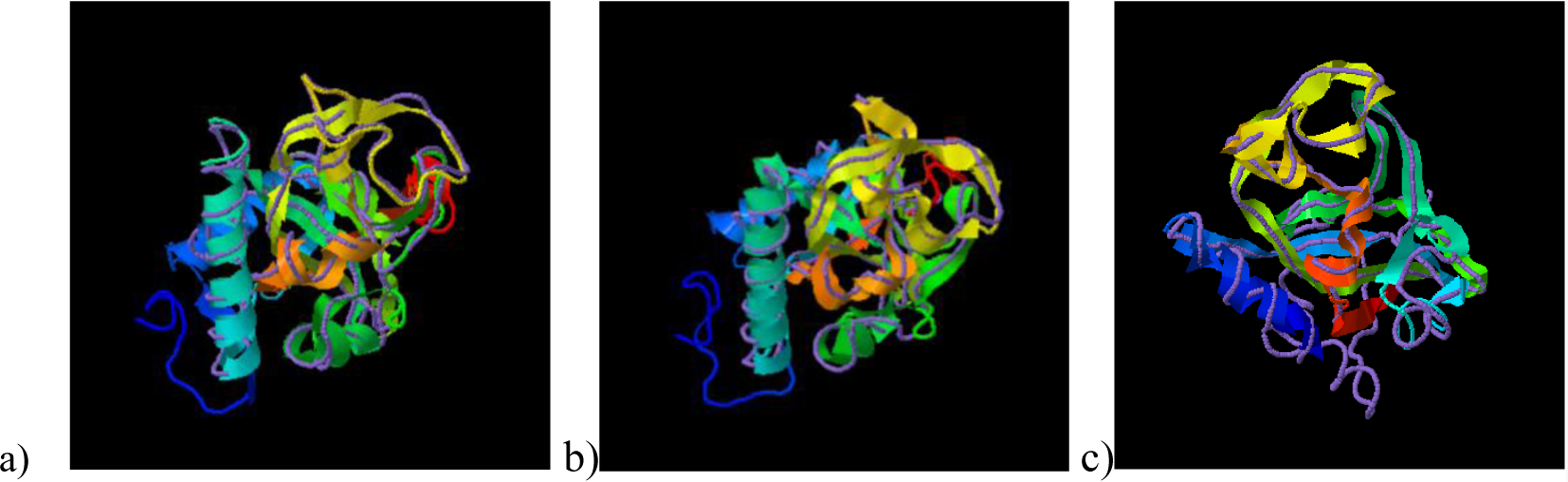
Superimposition of the PARP models of newly found Brassica homologues of RCD1 with the crystal structure of *A. thaliana* PARP (5ngoA) using TM-Align, where ribbon indicates Brassica PARP and discreet line indicates *A. thaliana* PARP. a) *B. carinata* b) *B. juncea* c) *B. oleracea*.

**Fig 4.**
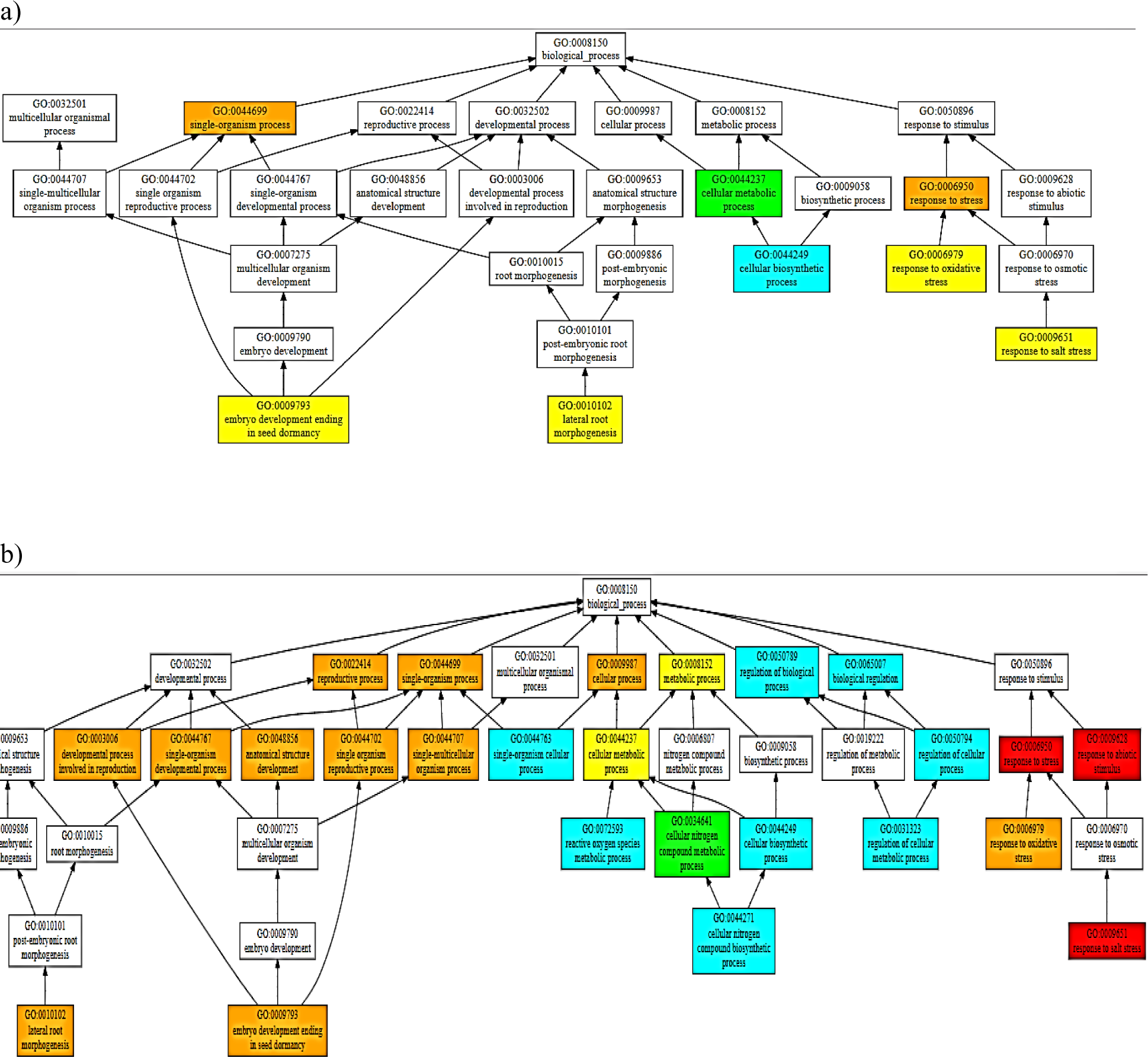
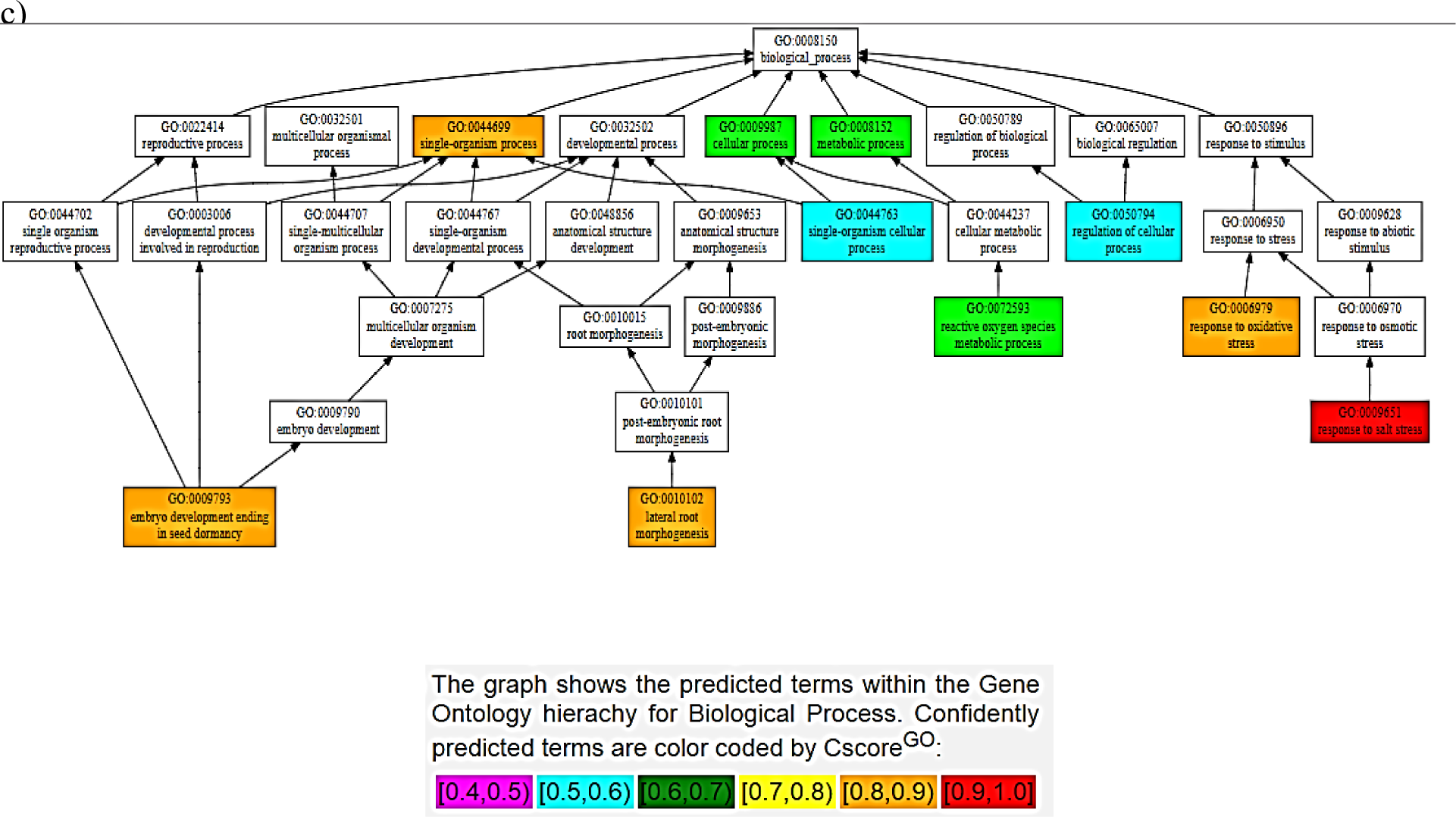
Consensus Prediction of Biological Function of the domains of BcRCD1, BoRCD1 and BjRCD1 a) WWE b) PARP c) RST

