## Supplementary material for "Identification of the Likely Orthologues of RCD1 within the Plant Family Brassicaceae": Figure 1: Folded and Unfolded areas within the Brassica Homologues of RCD1 a) B. carinata, b) B. juncea c) B. oleracea

Supplementary data:

a) 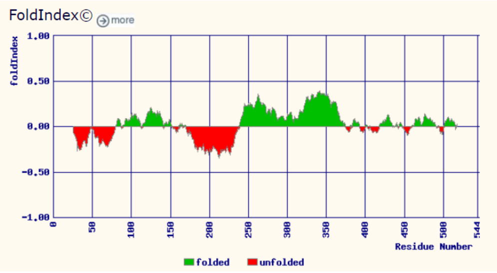


b) 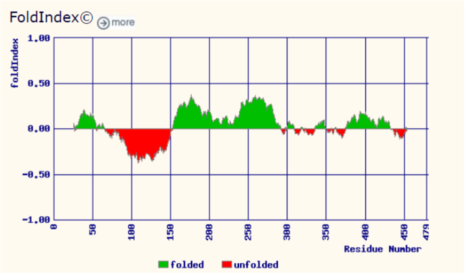


c) 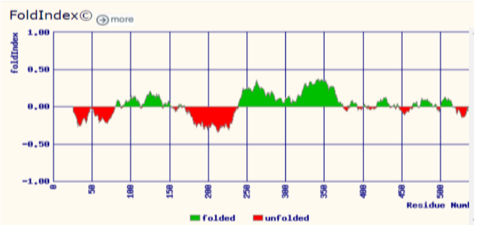


Figure 1: Folded and Unfolded areas within the Brassica Homologues of RCD1 a) *B. carinata*, b) *B. juncea* c) *B. oleracea*
